# Hygiene Hampers Competitive Release of Resistant Bacteria in the Commensal Microbiota

**DOI:** 10.1101/639443

**Authors:** Magnus Aspenberg, Sara Maad Sasane, Fredrik Nilsson, Sam P. Brown, Kristofer Wollein Waldetoft

**Affiliations:** Centre for Mathematical Sciences, Lund University, Box 118, 221 00 Lund, Sweden; Department of Clinical Pharmacology, 221 85 Lund University Hospital, Sweden; School of Biological Sciences, Georgia Institute of Technology, 311 Ferst Drive, Atlanta, GA, 30332 USA

**Keywords:** Antibiotic resistance, hygiene, competitive release, ecology, metacommunity ecology

## Abstract

Good hygiene, in both health care and the community, is central to containing the rise of antibiotic resistance, as well as to infection control more generally. But despite the well-known importance, the ecological mechanisms by which hygiene affects resistance evolution remain obscure. Using metacommunity ecology theory, we here propose that hygiene attenuates the effect of antibiotic selection pressure. Specifically, we predict that hygiene limits the scope for antibiotics to induce competitive release of resistant bacteria within treated hosts, and that this is due to a modulating effect of hygiene on the distribution of resistant and sensitive strains in the host population. We show this in a mathematical model of bacterial metacommunity dynamics, and test the results against data on antibiotic resistance, antibiotic treatment, and the use of alcohol-based hand rub in long-term care facilities. Our results underscore the importance of hygiene, and point to a concrete way to weaken the link between antibiotic use and increasing resistance.

## Introduction

Antibiotics have revolutionised modern medicine, but they also drive the evolution of resistance, and thereby contribute to their own demise (1, 2). To meet this challenge, there is intense work in both evolutionary theory and clinical practice to limit or optimise antibiotic use (3–5). A key task is to manage collateral antibiotic exposure of the commensal microbiota, of which medically important bacteria are often a part. Indeed, a recent analysis found that such ‘bystander selection’ dominates antibiotic exposure for common human pathogens (6).

In addition to the role of antibiotic use, there is within the medical profession a firm appreciation of the importance of hygiene in the management of antibiotic resistance. Measures of hygiene, such as hand disinfection, are fundamental to good medical practice after the discoveries of Semmelweis Ignác Fülöp in the mid 1800s (7), and form an integral part of current efforts to contain antibiotic resistant bacteria (8). Moreover, general hygiene and sanitation in the community are considered important in slowing the evolution of resistance, with the rationale that they limit antibiotic consumption by decreasing the incidence of infection (9).

It is thus recognised that antibiotic use and hygiene are key factors in the evolution of resistance, and that this evolution often takes place in a complex microbial community – but these insights have yet to be put together. This is the task of the present investigation: to study the joint effects of hygiene and antibiotic use on the rate of increase of resistant bacteria in microbial communities. It requires that we integrate the within-host ecological effects of antibiotics with the between-host process of bacterial dispersal, where the latter is affected by hygiene, and possibly other factors, such as population density or cultural habits. This sits most naturally within the theoretical framework of metacommunity ecology (10, 11), and we will frame both informal discussion and mathematical analysis using this approach. However, since metacommunity theory is new to antibiotic resistance studies, we also reproduce the key result within the epidemiological compartmental modelling framework that is currently standard in the field (12). (See Supplement A for details.)

The paper has three parts. First, we informally explain how metacommunity theory predicts that hygiene should limit the within-host ecological response to antibiotic treatment. Second, we develop a mathematical model of resistance dynamics in a metacommunity context, and show that this prediction obtains in the model. Third, we test the prediction against antibiotic resistance data from the European Centre for Disease Prevention and Control (ECDC), and find that it is consistent with the data.

## Metacommunity ecology implies that hygiene should attenuate competitive release

Bacteria form local communities (microbiotae; microbiomes) within individual hosts, and they transmit between host individuals. In ecological terms, bacteria thus form metacommunities, that is, networks of local communities interconnected by dispersal (13). Within each local community different bacterial strains compete with each other, such that the growth and abundance of a focal strain is limited by other strains. If the local community contains a mixture of antibiotic resistant and sensitive strains, antibiotic administration is expected to kill sensitive bacteria, and relieve resistant ones of competition, allowing them to proliferate. This is an ecological phenomenon known as competitive release (14–16). Previous work has indicated that it promotes the evolution of resistance, and that its strength depends on the dosage of the drug (17, 18).

However, the drug dose is only part of the picture. Competitive release of resistant bacteria requires that both sensitive and resistant strains are simultaneously present in the bacterial community within the treated host individual. And if they are, the magnitude of the release depends on the extent to which resistant strains are limited by competition from sensitive strains (see (16)). If resistant strains are absent within a host, they cannot increase (save for the possibility of *de novo* mutation). And, conversely, if they dominate the community already in the absence of antibiotics, they will gain little from the killing of the few sensitive cells that are present. In contrast, if resistant bacteria constitute but a small proportion of the competing community, the killing of the sensitive majority can decrease competition to a larger extent and result in a larger amplification of resistance (*Figure 1 A*).

**Figure 1.**
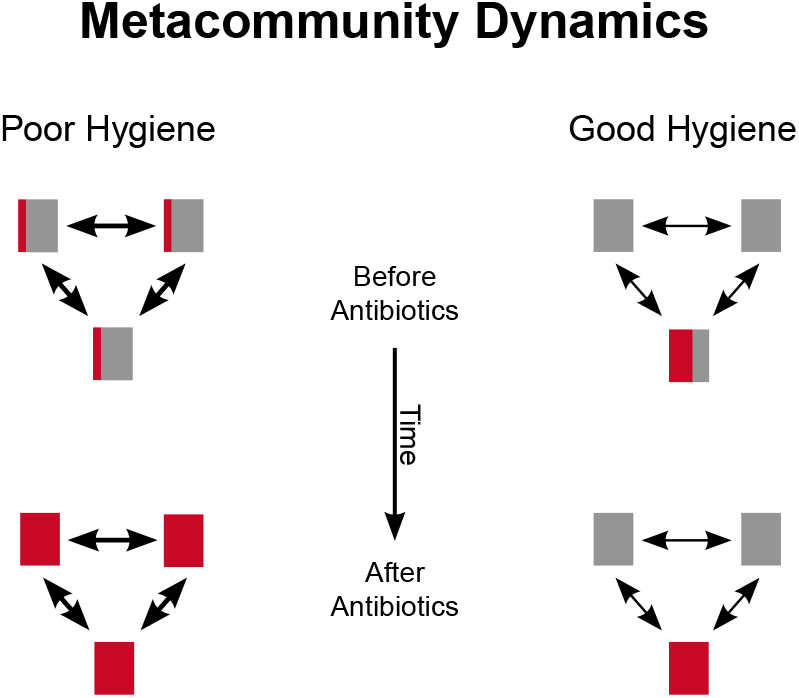
Schematic of metacommunity dynamics. Metacommunities with three host individuals are depicted. Before antibiotic treatment, resistant (red) and sensitive (grey) bacteria have similar total abundances in both poor and good hygiene settings, but they are differently distributed across host individuals. After antibiotic treatment and competitive release, resistant bacteria are more abundant in the poor hygiene setting. Red bars represent resistant strains, and grey bars sensitive strains, the area of each bar representing the abundance of the corresponding strain. Double arrows denote transmission of bacteria between hosts, thicker arrows indicating a higher rate of transmission.

As a consequence, the scope for competitive release is maximized when resistant strains are present in a large proportion of host individuals, and have a low abundance in the local communities in which they are present. Hence, for a given total abundance of resistant bacteria in the metacommunity, the competitive release will be larger when the resistant bacteria are more uniformly distributed across the host population.

The distribution of organisms in metacommunities is a focus of research in metacommunity ecology, and a picture has emerged that, perhaps unsurprisingly, dispersal of organisms among habitat patches tends to spread them across these patches, making them more uniformly distributed (13). Conversely, this means that interventions that prevent dispersal should make the distribution of organisms in the metacommunity less uniform than would otherwise have been the case, and tend to isolate different types of organism in different habitat patches.

In the context of humans carrying resistant and sensitive bacteria, measures of hygiene are precisely such interventions. They should therefore make the distribution of resistant strains less uniform, leaving them abundant in some individuals and absent from others, and thus decrease the scope for competitive release.

This leads to our prediction: Increasing the level of hygiene should decrease the effect of antibiotic pressure on the competitive release of resistant bacteria. That is to say that, whilst the magnitude of competitive release increases with antibiotic pressure, this increase is less steep when the level of hygiene is higher. We focus on hygiene, because it is readily actionable and its importance is well-established,but the prediction applies to any factor that limits bacterial dispersal among host individuals.

## Mathematical modelling of bacterial metacommunity dynamics supports the prediction

In the previous section, we informally explained why metacommunity ecology theory predicts that hygiene should attenuate competitive release. To test the validity of the argument, we now subject it to formal modelling. In the main text, we use a metacommunity model, and in Supplement A, we complement the analysis with an epidemiological compartmental model.

Our model of bacterial metacommunity dynamics is an extension of Hubbell’s neutral model of biogeography and relative species abundance (19, 20). The bacterial metacommunity is modelled as a set of interconnected bacterial communities in different host individuals. In these communities there are two types of bacteria – antibiotic resistant and antibiotic sensitive – and the total number of bacteria (resistant and sensitive) in each host individual is constant, as is the number of host individuals. These constants are denoted by *J* for the number of bacteria in each host and *N* for the number of host individuals. The constant community size means that the resistant and sensitive types compete with each other, and a decrease in one entails an equal increase in the other, representing competitive release.

The model tracks the proportion of bacterial cells that are resistant in the metacommunity, as time, step by step, unfolds. In each time step bacterial cells die. In the absence of antibiotics, one cell dies in each host individual, and the resistance status of a cell does not affect its probability of dying. With increasing antibiotic pressure, however, more cells can die in each time step, and the death process becomes increasingly biased towards sensitive cells (see Supplement A for details). We expand on this below.

Once those cells have died, they are replaced by new cells by division. These cells may originate from within the same host or from different host individuals. The probability that a new cell comes from a different host depends on the level of hygiene – the better the hygiene, the lower the probability that a bacterium transmits from another host. If the new cell originates from a different host, the probability that it is resistant is equal to the proportion of cells that are resistant in the metacommunity as a whole, and if it originates from within the focal host, the probability of resistance equals the proportion of cells that are resistant in that host.

Let us compute the probabilities more precisely. Let *R_j_* (*t*) be the number of resistant bacteria at a given time *t* in host *j*. Since we are only looking at the transition between two adjacent time steps, we will drop the index *t* and only write *R_j_*. Each time step has two phases. In the *first phase* one or several bacteria are lost, and in the *second phase* these bacteria are replaced. In the first phase, given *R_j_* number of resistant bacteria in an individual, the probability that a resistant bacterium dies is

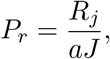

where *a* ≥ 1 is the antibiotic pressure, represented as the differential survival of resistant and sensitive cells. Hence, *a* = 1 means that no antibiotics are present, and the probability that it is a resistant (or a sensitive) cell that dies is simply proportional to their number, whereas *a* > 1 means that antibiotic treatment makes the death process biased towards sensitive cells.

The probability that sensitive bacteria are lost is

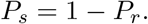

The number of sensitive bacteria lost is an integer function *ρ*(*R*, *a*) that depends on the antibiotic pressure *a* and the abundance *R* = *R_j_* of resistant bacteria in host *j* (or equivalently, the abundance of sensitive bacteria *J* – *R*). However, in the computations (see Supplement A), we consider *ρ* to be a continuous function that is differentiable in the *a*-direction. Moreover, we require that it is a decreasing function of *R*, meaning that more sensitive bacteria die if there are many such bacteria. We then take the integer part of this function to go back to the integer valued *ρ*. Let us denote by *μ*(*R*, *a*) the number of bacteria lost in this first phase (*i.e.*, *μ*(*R*, *a*) = 1 or *μ*(*R*, *a*) = *ρ*(*R*, *a*)).

In the second phase, given *R* number of resistant bacteria, the probability that one resistant or one sensitive bacterium reappears is, respectively,

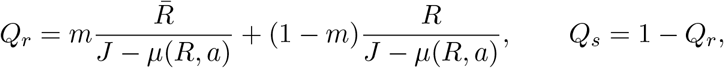

where

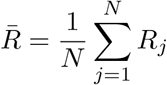

is the average number of resistant bacteria in the whole population at that time, and *m* ∈ [0,1] (called the migration rate in the literature) is the probability that a bacterium is chosen from the metacommunity as a whole, rather than the focal individual. The hygiene parameter *h* = 1 – *m*, so that good hygiene means high *h*-values and low *m*-values (and vice versa). This procedure is performed precisely *μ*(*R*, *a*) times to replace all bacteria lost in the first phase.

However, the time steps can be made very small in the mathematical model, and therefore we assume that the portion *μ*(*R*, *a*)/*J* is very small, although the number *μ*(*R*, *a*) can be large (in particular much larger than 1). This means that we may replace 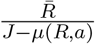 by 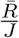 (with a very small error). The new probabilities *Q_r_* and *Q_s_* then become

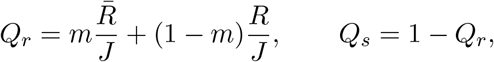

We compute the expected change of the number of resistant bacteria in one time step. This expectation is denoted by *Z* = *Z*(*a*, *h*), which is a function of *a* and *h*. We prove that

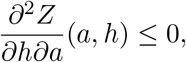

if *a* ≥ 1, with strict inequality if *a* > 1. Hence, the effect of antibiotic pressure on the rate of increase of resistance (*∂Z*/*∂a*) decreases with improving hygiene.

We also show that this is due to the effect of hygiene on the *β*-diversity of the metacommunity. (See Supplement A for details.) The *β*-diversity is the standard deviation of the number of resistant bacteria *R_j_* across host individuals.

## Resistance data are consistent with the prediction

In the preceding sections, we saw, first, that informal theory predicts that hygiene attenuates the effect of antibiotics on the competitive release – and thus the rate of increase – of resistant bacteria, and, second, that formal modelling supports thevalidity of that argument. Now, we test this prediction empirically. To do so, we restate it in statistical form.

The prediction is as follows: If a measure of the increase of resistant bacteria in a given setting is regressed upon a measure of antibiotic pressure (*a*) and a measure of hygiene (*h*) in that setting, there should be an interaction (*a* · *h*), and this should be negative. The reason is that the interaction term represents the effect that the value of one variable (e.g., *h*) has on the effect of the other variable (e.g., *a*), a negative term meaning attenuation.

We tested this prediction against data on resistance to third generation cephalosporins in *Enterobacteriaceae,* the use of *β*-lactam antibiotics, and the use of alcohol-based hand rub in long-term care facilities in different European countries, reported by the European Centre for Disease Prevention and Control (ECDC). We chose these data because, as far as we are aware, this is the only data set that is large enough, and contains information on antibiotic resistance, antibiotic use, and a reliable measure of hygiene, all of which are necessary to test the prediction. The principal shortcoming of the data is that they are cross sectional, and thus only provide the level of resistance at a given time, not the rate of increase. We therefore modelled the increase in resistance as the enrichment of resistant strains in the long-term care facilities in each country as compared to resistance in *E. coli* in the general society in that country. (See Supplement B for details on the data and analysis.)

The data were analysed by logistic regression (using R ver. 3.5.0, see (21)). In accordance with the prediction, there was a significant negative interaction between hygiene and antibiotic pressure (*p* = 0.005), and this amounted to a pronounced effect of hygiene on the slope of resistance on antibiotic pressure. The data and the regression model are shown in *Figure 2 A*, and the change in the relationship between antibiotic pressure and antibiotic resistance associated with an increase from medium-low to medium-high consumption of hand rub is illustrated in *Figure 2 B*.

**Figure 2.**
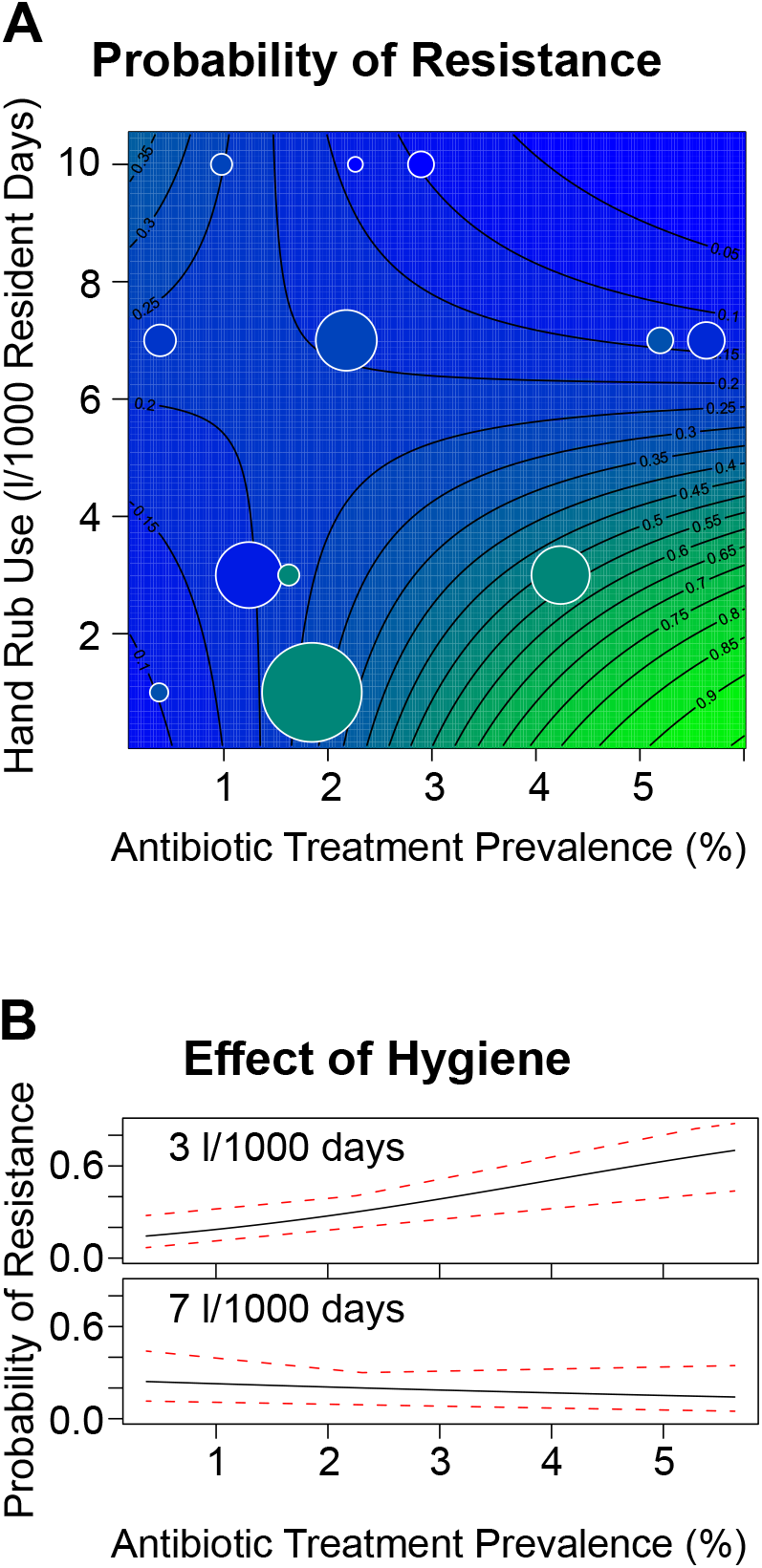
**A. Resistance data and statistical model.** The ECDC resistance data are overlain a contour plot of the logistic regression model that was fitted to them. The probability that an *Enterobacteriaceae* isolate is resistant to third generation cephalosporins is plotted against the consumption of alcohol-based hand rub (vertical) and the point prevalence of *β*-lactam treatment (horizontal). The colour and contours represent the probability of resistance. Bubbles represent data reported for individual countries, the area being scaled to the number of bacterial isolates tested. (See Supplement B for details.) **B. The effect of a moderate improvement in hygiene.** Based on the data and analysis given in (B), the probability that an isolate is resistant (vertical) is plotted against the prevalence of **β**-lactam treatment (horizontal). The upper and lower panels show this relationship for a hand rub consumption of 3 and 7 litres, respectively, per 1000 resident days. Red dotted lines represent 95 percent confidence bands. (See Supplement B for details.)

## Discussion

Here we have employed metacommunity ecology theory to study what are arguably the two most important levers available to limit the rise of resistance – the amount of antibiotics used and the level of hygiene. We found that informal theory, formal modelling, and data from the ECDC all yield the same conclusion: Improvements in hygiene weaken the link between antibiotic use and increasing resistance. The mechanism is that hygiene limits bacterial dispersal, and thereby alters the diversity that is the substrate for antibiotic induced competitive release. Since a quantitative assessment of this effect in the mathematical model requires biological information that is currently unavailable, we assessed the effect size directly with the ECDC resistance data, and found that it was quite striking (*Figure 2 B*).

Naturally, there are caveats to the conclusion. The data set is relatively small and pertains to a particular setting – long-term care facilities. In addition, the frequency of resistance in these facilities as compared to the baseline frequency in each country, though a measure of enrichment of resistant bacteria, is not strictly a rate of increase. Furthermore, the results may be confounded by factors not included in the analysis. For example, since each data point represents a different country, there may be differences in health care systems or cultural habits that affect the outcome.

On the other hand, the validity of the argument is supported by the fact that a simple compartmental model – which is mathematically very different from the metacommunity model – whilst unable to capture the nuances of bacterial diversity, does indicate the same qualitative conclusion. (See Supplement A for details.) In addition, there are previous empirical results that are suggestive of a role for bacterial dispersal in modulating the effect of antibiotic pressure. Bruinsma *et al.* (22) investigated the resistance to several antibiotics in *E. coli* and *Enterococci* in the faeces of healthy volunteers living in cities with different population densities, and found that it correlated poorly with the antibiotic pressure per individual, but well with the antibiotic pressure per land area. Assuming that bacterial dispersal among individuals is facilitated by a high population density, this fits well with a modulating effect of dispersal, since the antibiotic pressure per land area is the product of the pressure per individual and the population density, and thus corresponds to the interaction term, *a* · *h*, in the statistical model above.

Whilst decreasing the population density may not be a viable route to limiting antibiotic resistance, improving health care hygiene is. Proper hand hygiene in clinical work is a cornerstone of patient safety (23), but more than 150 years after Semmelweis compliance is still poor (24). This is unfortunate, because hand hygiene stands out among possible anti-resistance interventions in that it is simple, safe, and cheap, and should therefore be possible to implement rapidly, and without major issues. Furthermore, as suggested by the findings of Bruinsma *et al.* (22) above, the effect of dispersal limitation on competitive release should not be confined to the health care setting, but apply to the general community as well.

In conclusion, we have here introduced metacommunity ecology theory to the study of antibiotic resistance, and found that interventions to limit microbial dispersal provide a means to attenuate the effect of antibiotic selection pressure on the rise of resistance.

## Acknowledgments

The authors wish to thank Luke McNally, Erik Wahlén, Tomas Persson, Tatiana Turova, Lars Råberg, Rolf Lood, James Gurney, Yifei Wang, the Brown lab, and the biology and mathematics seminar in Lund for their advice at various stages of the project.

This work was supported by grants from the Wenner-Gren Foundations, the Royal Physiographic Society of Lund (the Fund of the Hedda and John Forssman Foundation), the Sten K Johnsson Foundation, the Crafoord Foundation, and the Centers for Disease Control and Prevention (BAA 2017-0ADS-01).

## Supplement A: The mathematical models

### Analysis of the metacommunity model

#### 1. Introduction

The mathematical model of the interaction between antibiotic pressure and hygiene is based on an idea by S. Hubbell (20). Let us here explain the basic mechanics in our model. Let *N* be the number of individuals in the host population, *J* the total number of resistant and sensitive bacteria in each individual host, *m* a hygiene parameter, and *a* a measure of the antibiotic pressure. In each time step, there are two phases. In the first phase, each host individual either loses a resistant bacterium or a number *ρ* ≥ 1 of sensitive bacteria. In the second phase the individual gains the same number of bacteria of possibly different types (sensitive or resistant). Hence, after each time step, the total number of bacteria is unchanged. Let *R* be the number of resistant bacteria at some given time for a given host individual, and 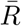 the average portion of resistant bacteria over the whole metacommunity; *i.e*.,

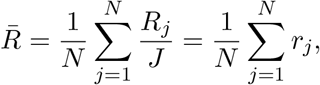

where *R_j_* is the number of resistant bacteria for host *j* and *r_j_* = *R_j_*/*J*. We then have the possibilities that the change of the number of resistant bacteria for an individual can attain the values {− 1, 0,…, *ρ*} after these two phases have taken place.

The probability that a resistant bacterium is lost, given *R* = *R_j_* number of resistant bacteria in host *j*, is

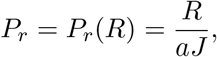

where *a* is the antibiotic pressure (*a* = 1 means that there are no antibiotics). The probability that *ρ* sensitive bacteria are lost is

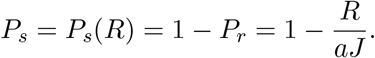

Let us briefly describe the properties of *ρ* = *ρ*(*x*, *a*), which is a function of the antibiotic pressure *a* and the abundance *x* of resistant bacteria (or equivalently, the abundance of sensitive bacteria *J* − *x*). We may also consider *ρ* as a function of the relative abundance of *x* or *J* − *x* since the total number of bacteria in each host is constant after each time step. We require that *ρ* is decreasing in *x*, meaning that more sensitive bacteria die if there are many such bacteria. We will also need a technical condition on *ρ* including its *a*-derivative (written in the Proposition below). This condition is discussed after the Proposition, and there are many “natural” functions *ρ* that satisfy it (for instance *ρ* must increase with higher *a*-values). Let us denote by *μ*(*x*, *a*) the number of bacteria lost in this first phase (*i.e*. *μ*(*x*, *a*) = 1 or *μ*(*x*, *a*) = *ρ*(*x*, *a*)).

Let us now turn to the second phase. Using Hubbell’s model with some modifications (Hubbell does not include the *a*-variable), the probabilities that a resistant or a sensitive bacterium reappears are, respectively,

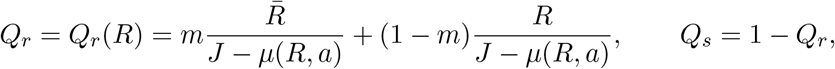

where

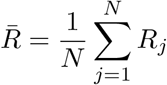

is the average number of resistant bacteria in the metacommunity, *R* = *R_j_* for some *j*, and *m* ∈ [0, 1] is the probability that a bacterium is chosen from the metacommunity as a whole, rather than from the focal individual.

However, since each time step (the two phases) can be arbitrarily small, at least in the mathematical model, we will in this model make the assumption that *μ*(*R*, *a*)/*J* is very small (although the number *μ*(*R*, *a*) can still be large), so that 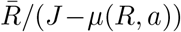 is very close to 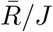. With this approximation, the probability that one resistant or one sensitive bacterium reappears is, again given *R* number of resistant bacteria, respectively,

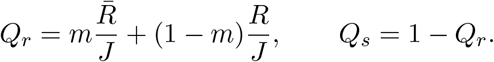

We call *h* = 1 − *m the hygiene parameter*, so that good hygiene means high *h*-values and low *m*-values (and vice versa). This procedure is performed precisely *μ*(*R*, *a*) times to replace all bacteria lost in the first phase.

If we look at the transition probabilities for all hosts, this creates a huge Markov chain, since the number *N* of people and the total number *J* of bacteria are usually very large numbers. We will instead consider the expected change *Z* of resistant bacteria in the whole population, given an antibiotic pressure *a*, the level of hygiene *h*, and the number of resistant bacteria *R_j_* in each host *j*, 1 ≤ *j* ≤ *N*. So *Z* is a function of *a*, *h*, and the numbers *R_j_*, 1 ≤ *j* ≤ *N*.

In the proposition below, we note first that the function *ρ*(*x*, *a*) is in reality integer valued. However, in the proposition it is assumed to satisfy a Lipschitz continuity condition (with differentiability in a). We then take the integer part to go back to its real meaning. When one takes the integer part of *ρ*, the partial derivatives in the proposition have to be replaced by corresponding “discrete” derivatives.

On the other hand, one can view the death and rebirth processes of bacteria as binomially distributed; the death process *X* ~ *Bin*(*ρ*/*J*, *J*) and the rebirth process *Y*, where *Y*|*X* ~ *Bin*(*Q_r_*, *X*) and hence 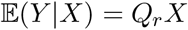 is the conditional expectation. Then, since 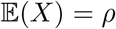, we have

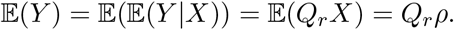

As *J* → ∞, we note that the death binomial process will converge to a Poisson process.

In the proposition below, *ρ*(*x*, *a*) is considered as a function of the relative abundance *x* of bacteria, *i.e*., *x* ∈ [0, 1]. Write 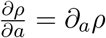.

##### Proposition 1.1.

*Given Z* = *Z*(*a*, *h*) *as above. Suppose that the function ρ*(*x*, *a*) ≥ 1 *is decreasing in x, ρ*(*x*, *a*) *and ∂_a_ρ*(*x*, *a*) ≥ 0 *are Lipschitz continuous, ρ*(*x*, 1) = 1 *and that*

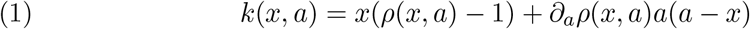

*is decreasing in x. Then*,

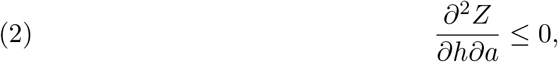

*with strict inequality if a* > 1.

In the computations we will use the parameter *m* = 1 − *h* and hence prove the equivalent statement

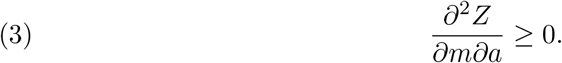

We call this *the first prediction*.

We now discuss the condition (1). If for example *ρ*(*x*, *a*) = *ϕ*(*a*)*h*(*x*) + 1, where *h*(*x*) is decreasing in *x* then (1) is equivalent to the statement that

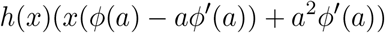

is decreasing in *x*. Since *k*(*x*, *a*) ≥ 0, this is true in particular if *ϕ*(*a*) − *aϕ′*(*a*) ≤ 0 which is equivalent to

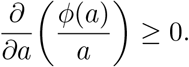

There are many functions that satisfy this condition (1). A simple example is *ρ*(*x*, *a*) = 1 + (*a* − 1)(1 − *x*). We will also write 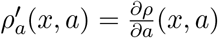.

As a by–product we have:

*The greater the β–diversity of the metacommunity, the smaller the competitive release of resistant bacteria in response to antibiotic pressure*.

We call this the *second prediction*.

#### 2. Proof of Proposition 1.1

The aim in this section is to study the expected change in the number of resistant bacteria in the whole population, given certain values on *a* (antibiotic pressure) and *m* (hygiene). Also, we will see that there is an interplay between the *β*–diversity of the metacommunity and its sensitivity to antibiotic pressure.

In the first phase, a number of bacteria are lost; one resistant with probability *P_r_* or *ρ* sensitive with probability *P_s_* = 1 −*P_r_*. In order to have a zero sum process, all bacteria lost in the first phase must be replaced by new ones in the second phase. So if *ρ* sensitive bacteria were lost in the first phase (in some host), then obviously *ρ* new bacteria must be reborn, and if a resistant bacterium was lost in the first phase, we only need to add one extra bacterium in the second phase.

We proceed as in Hubbell’s model, but perform the bacterial replacement procedure *ρ* times if sensitive bacteria were lost in the first phase, and only once if a resistant bacterium was lost in the first phase. Recall *Q_r_* = *Q_r_*(*R_j_*) and *Q_s_* = *Q_s_*(*R_j_*) for the probabilities for host *j* that one resistant or sensitive bacterium reappears, respectively.

We let *X* be the total number of resistant bacteria lost in the first phase and *Y* be the total number of resistant bacteria that reappear in the second phase. Moreover, let *X_j_* be the number of resistant bacteria lost for host individual *j* in the first phase, and *Y_j_* the number of reappearing resistant bacteria for host individual *j* in the second phase. So we have

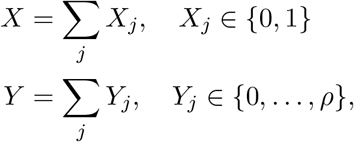

where *ρ* = *ρ*(*R_j_*, *a*). We then put

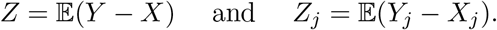

Then *Z* is the expected total change of resistant bacteria after one full time step (both phase one and phase two) and

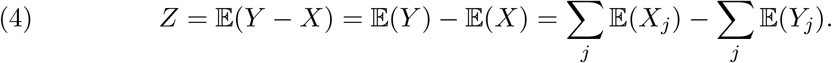

To compute 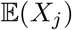 is easy:

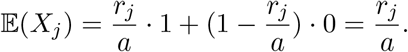

To compute 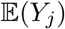 is more complicated since they depend on *X_k_*, 1 ≤ *k* ≤ *N*. If a resistant bacterium was lost in the first phase (*i.e*., *X_j_* = 1), then the expectation of the change of resistant bacteria in phase two is (recall the probabilities *Q_r_* for picking a resistant and 1 − *Q_r_* for a sensitive bacterium)

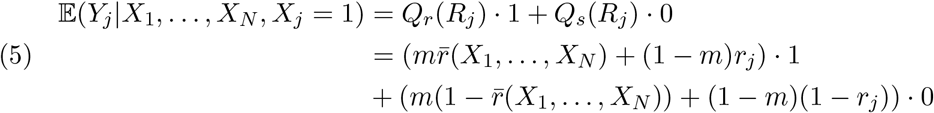

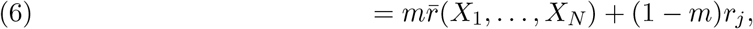

where 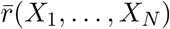 is the average of all *r_j_ after* the first phase (and *X_j_* = 1). If sensitive bacteria were lost in the first phase then perform the growth process as above but *ρ*(*R_j_*, *a*) times and consequently

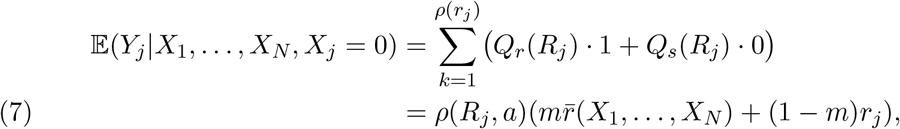

where *X_j_* = 0. Now, 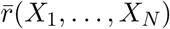 can change quite drastically if a lot of sensitive bacteria were lost in the first phase, but not if resistant bacteria were lost in the first phase. However, we assume that the portion of resistant bacteria does not change too much after the first phase, so we just use 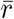 and *r_j_* as they were *before* the first phase. This is equivalent to saying that the quotient *ρ/J* is very small, which is the assumption made in the previous section (so that *Q_r_* does not depend on *μ*(*R*, *a*)). However, the outcome of *Y_j_* still depends drastically on *X_j_*. Let us write *ρ*(*R_j_*, *a*) = *ρ*(*r_j_*, *a*) to simplify the notation in the coming computations, so that we regard *ρ*(*r_j_*, *a*) also as a function of the relative abundance of resistant bacteria for individual *j*. We have

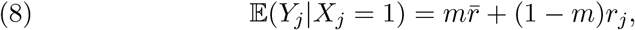

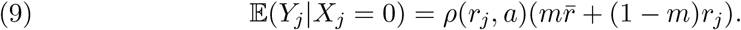

We now get

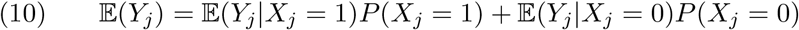

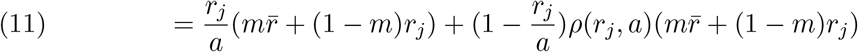

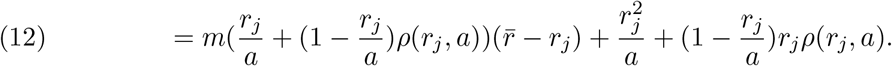

Consequently,

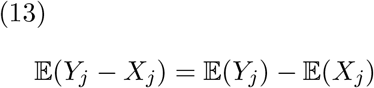

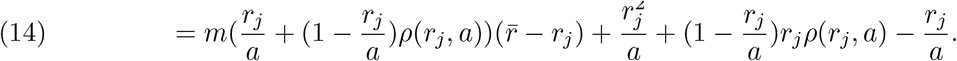

So

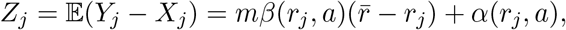

where

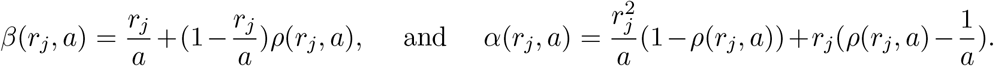

An easy computation yields 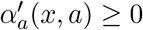 and *α* > 0.

We want to show that *∂Z_j_*/*∂a* increases in *m*, *i.e*., that *∂*^2^*Z_j_*/*∂m∂a* ≥ 0. To do this, we will first need that 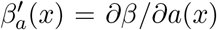 is positive and decreasing in *x*. We have

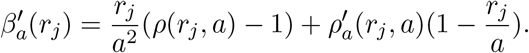

Since 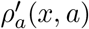 is positive from the assumptions and *ρ*(*r_j_, a*) ≥ 1, we clearly have *β′*(*a*) ≥ 0. We may without loss of generality order the *r_j_* so that *r*_1_ ≤ *r*_2_ ≤ … ≤ *r_N_*. Recall that both *ρ* and 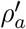 are decreasing functions in *x*. The condition (1) means that 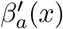 is decreasing in *x* and hence 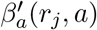 is decreasing in *j*. To prove (3), we need the following lemma, which we state in a more general form.

##### Lemma 2.1.

*Let ξ* ≥ 0 *be a decreasing function and f* ≥ 0, *both defined on a compact interval I. Suppose that μ*(*I*) = 1. *Then*

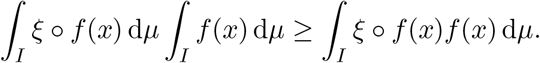

T. Persson pointed out that this lemma is quite similar to the Chebyshev integral inequality and he supplied an elegant proof based on this. However, for our purposes we take another route.

*Proof*. Let *α*(*x*) = *f*(*x*) − ∫*f*(*x*) d*μ*. Then ∫*α*(*x*) d*μ* = 0. Put *A*={*x*: *α*(*x*) ≥ 0} and *B*={*x*: *α*(*x*) < 0}. Since *ξ*(*x*) is decreasing, we have

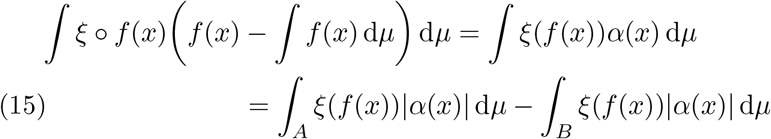

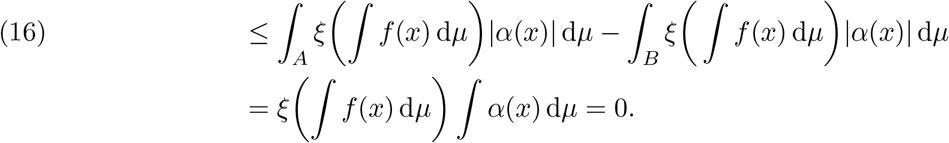

This proves the lemma.

##### Remark 2.2.

It also follows from the proof of the lemma that there are constants *c, c′* ≥ 0 which only depend on *ξ* such that

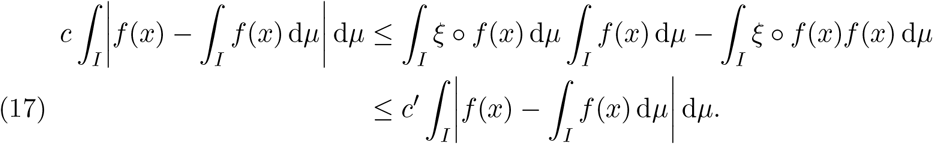

Indeed, the difference between the first terms on the lines (15) and (16) is

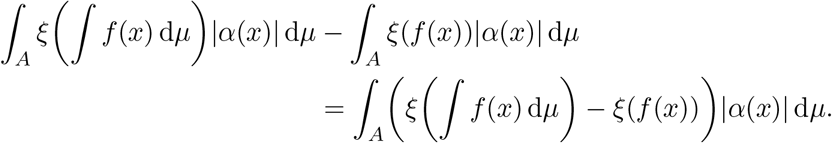

Since *ξ* is Lipschitz continuous, and since *α*(*x*) ≥ 0 on *A*, we conclude that there are constants *c*_1_ ≥ 0 and *c*_2_ ≥ 0 that only depend on *ξ* such that

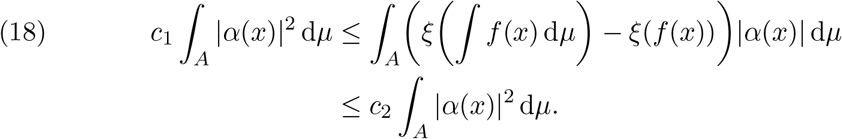

A similar inequality holds for the difference of the two last terms in (15) and (16). Indeed, by the same argument, remembering that *α*(*x*) ≤ 0 on *B*, there are constants *d*_1_ ≥ 0 and *d*_2_ ≥ 0 such that

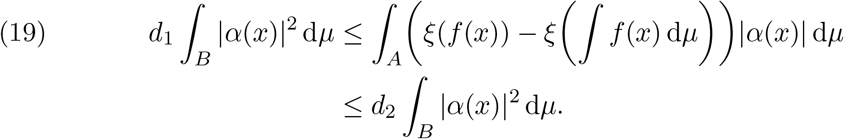

So we see that

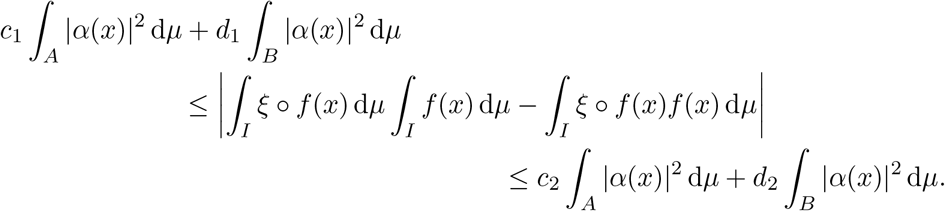

Taking *c* = min(*c*_1_, *d*_1_) and *c′* = max(*c*_2_, *d*_2_) gives the desired result.

Recall that 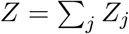 can be written as

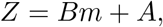

where

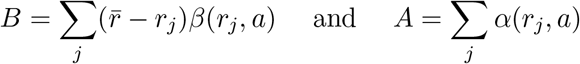

both depend on *a*. Therefore, with 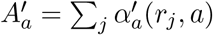, we haveand

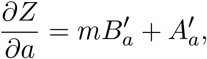

and

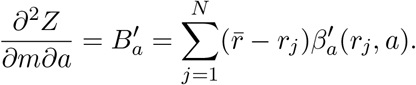

Applying the lemma to the uniformly distributed point measure 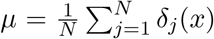 on the interval *I* = [1, *N*], where *f* (*j*) = *r_j_* and 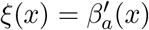 it follows that

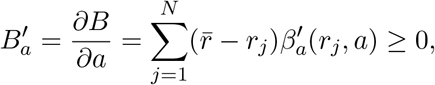

with strict inequality if *a* > 1 and the distribution of *r_j_* is such that not all are equal. In particular, if the *β*-diversity is zero, *i.e.,* the distribution of the resistant bacteria is completely uniform, then 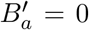. This proves our first prediction (Proposition 1.1).

We also note that *∂Z/∂a* is clearly positive since every 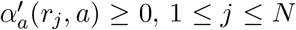 (this is simply equivalent to the unsurprising fact that the greater the antibiotic use in the population, the faster the proportion of bacteria that are resistant increases).

From the remark after the lemma, putting 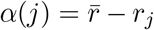, we see that the sum 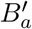 is comparable to the deviation:

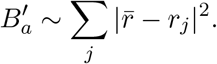

Hence, we have also settled the second prediction.

##### Remark 2.3.

A natural question is to ask whether

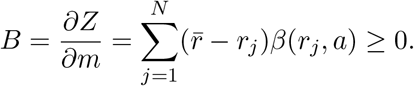

But this follows easily from the fact that *∂*^2^*Z/∂m∂a* ≥ 0 and integrating. Suppose that *m*_1_ ≤ *m*_2_ and fix some *a* ≥ 1. Then

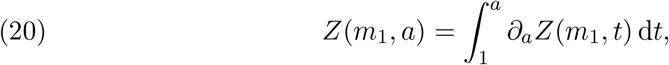

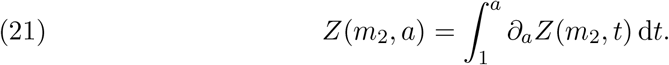

Since *∂_a_Z* is increasing in *m*, we have *Z*(*m*_1_, *a*) ≤ *Z*(*m*_2_,*a*) for all *a* ≥ 1 and all choices of *m*_1_ ≤ *m*_2_. Hence, *B* ≥ 0.

##### Remark 2.4.

Another consequence of the model is that if no antibiotics are present, then *Z* will no longer depend on *m*. Indeed, since *ρ*(*x*, 1) = 1, we have

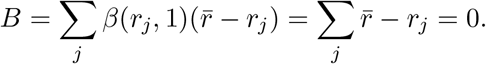

Hence, hygiene has no effect on the rate of increase of the number of resistant bacteria in the absence of antibiotics.

### Analysis of a simple compartmental model

It is common to study the spread of antibiotic resistant bacteria using the epidemiological compartmental modelling framework (12), and to connect our results to this rich tradition, we here show the main result of our study within this framework. The within-host diversity of bacteria is modelled as a subdivision of the host population into compartments with different portions of different bacterial strains. Since this is necessarily a crude representation of diversity, we will opt for simplicity rather than nuance in the choice of model. We begin with a model with 4 compartments, and then simplify.

Consider a commensal that is ubiquitous, or nearly so, such as *E. coli*. There are two strains, a sensitive strain and a resistant strain. And hosts are divided into compartments based on which strain(s) they harbour. Hosts in compartment *S* have only the sensitive strain, and those in *R* only the resistant. Hosts in compartment *S_R_* harbour mostly sensitive bacteria, but there are a few resistant cells, and hosts in *R_S_* have mostly resistant bacteria, but there are a few sensitive cells. Hosts flow from *S* to *S_R_* as they contract resistant bacteria by transmission. From *S_R_* they can either return to *S*, as the sensitive bacteria outcompete the resistant bacteria due to a fitness cost of resistance, or continue to *R* due to competitive release under antibiotic treatment that kills the sensitive cells. From *R*, hosts flow to *R_S_* by transmission of sensitive bacteria, and from there either return to *R* due to treatment, or transition to *S* due to a cost of resistance. The model is represented in *Figure S1 A*.

For simplicity, now assume that the cost of resistance is negligible, such that there are no flows from *S_R_* to *S* or from *R_S_* to *S*. The resulting model is represented in *Figure S1 B*. Since hosts in *R* and *RS* cannot leave this pair of compartments, and *R_S_* is dominated by resistant bacteria, *R* and *R_S_* can be combined. This yields the model represented in *Figure S1 C*. This model is described by the following ordinary differential equations:

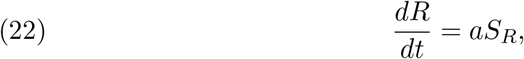

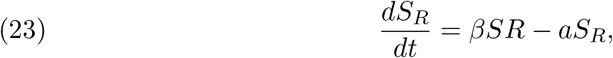

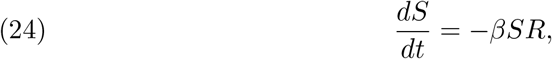

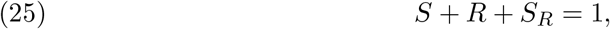

where *a* is the antibiotic pressure on the host population and *β* is the transmission rate. Improvements in hygiene lower *β*. Competitive release under antibiotic treatment is represented by the flow from *S_R_* to *R* according to *aS_R_.*

Let us first discuss the model informally to build intuition. The first prediction, that hygiene should attenuate the effect of antibiotic pressure on the competitive release of resistant bacteria, can be intuited by noting, firstly, that the rate at which *R* increases in the model in *Figure S1 C* depends on both the antibiotic pressure, *a,* and the proportion of hosts that are in compartment *S_R_,* and, secondly, that the rate at which new hosts enter compartment *S*_R_, and thus become available to treatment induced competitive release, depends on *β*, which, in turn, decreases with hygiene. The second prediction, that this is because improvements in hygiene make resistant strains less uniformly distributed across the host population, can also be intuited from the model. Note, firstly, that the distribution of resistant strains is most uniform when *S_R_* is large, and, secondly, that the rate of increase in *R* depends on Sr. Put another way, a large *S_R_* corresponds to the situation discussed in the main text, where resistant bacteria are present in many host individuals, but have a low abundance in the hosts in which they are present.

**Figure 3.**
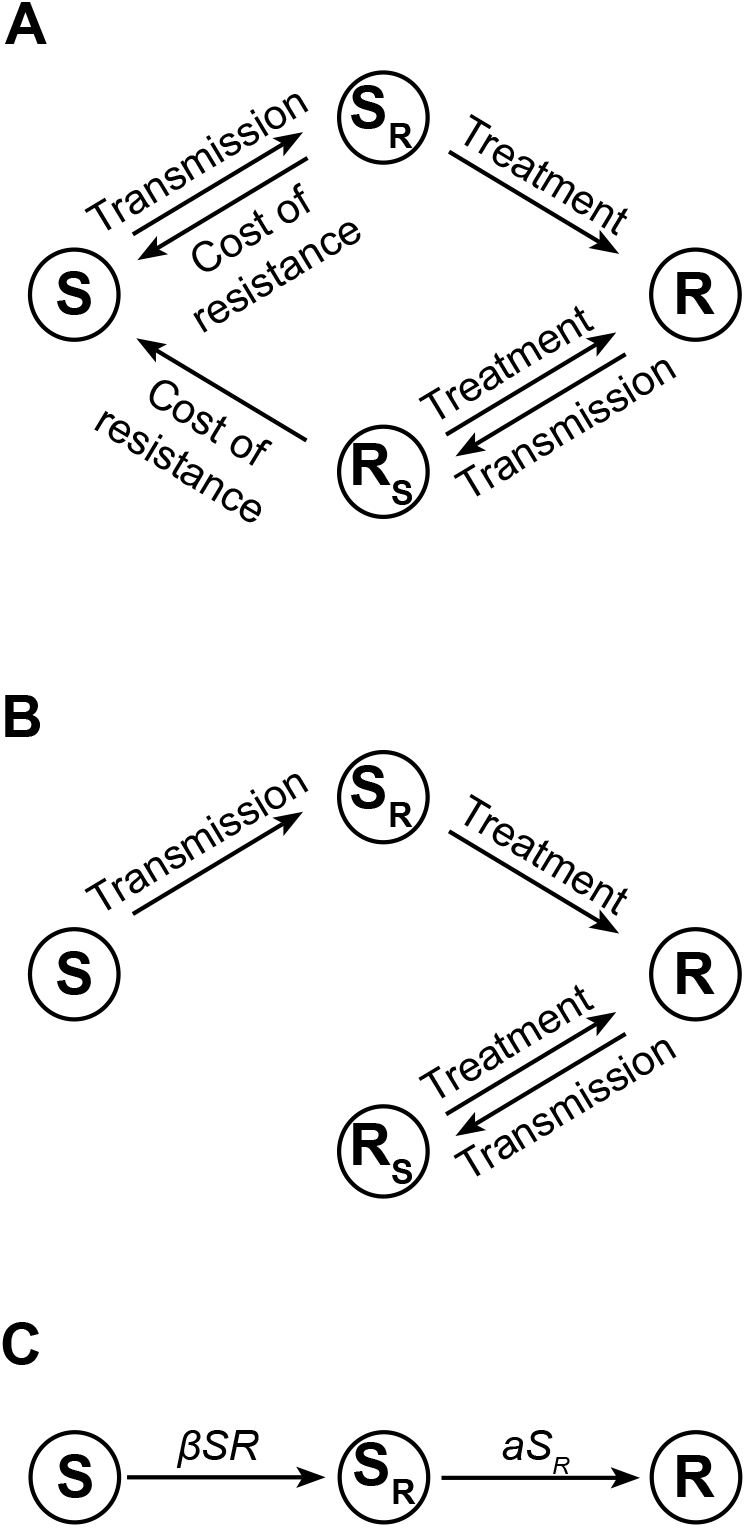

We now give a sketch of a proof that the first (and main) prediction follows from this model, at least under some circumstances (conditions on the prevalences of *R* and *S_R_*). We stress that this prediction holds *locally around any given initial values on R and S_R_*. With some conditions on *R* and *S_R_*, we can prove this globally as well, meaning far away from the initial values. These conditions can most likely be relaxed, at least up to some point, but for our purposes we only wish to illustrate that the same phenomenon can be found in this compartment model as in the metacommunity model.

Assume that the system of differential equations above is defined for *t* ≥ 0 with some initial values *S_R_*(0) and *R*(0). The first prediction is equivalent to the proposition that the second partial derivative of *R* satisfies

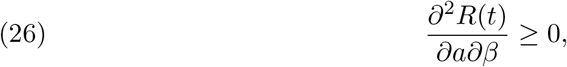

with strict inequality for *t* > 0. Locally, *i.e.,* for *t* close to 0 this is always true, as we will see below.

Let us first reduce the equations to a system in two variables, in *R* and *S_R_*. Let us also use the simpler variables *x* = *R* and *y* = *S_R_*:

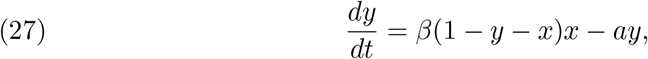

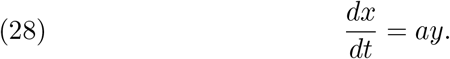

By making a simple change of variables (e.g. 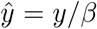, 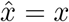) one can easily see that (26) is satisfied locally around any initial point *y*_0_ = *y*(*x*_0_), where *x*_0_ = *x*(0) and *y*_0_ = *y*(0), (note that the initial point is not dependent on the parameters *a* and *β*). However, in order to analyse the long term behaviour, we take another route.

Eliminating the *t*-variable, we get

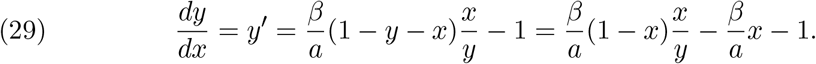

Hence, the solution curves *y* = *y*(*x*) only depend on 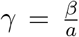 and the initial value *y*(*x*_0_) = *y*_0_. So let us keep in mind that *y* = *y*(*x*, *γ*), and write

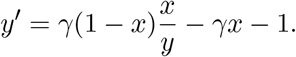

We also note that *R*′ = *x*′ = *ay*(*x, γ*), and thus, after some calculations,

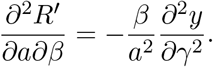

We recall that both *a* and *β* are positive numbers. So if we can show that 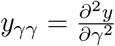 is negative, the result follows by integrating in the time-domain.

We first show that 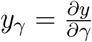 is non-negative. Differentiating (29) with respect to *γ* we get,

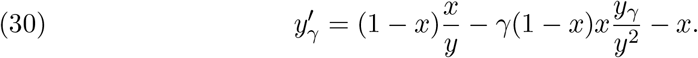

Considering this as a linear equation in *y_γ_*, assuming that a solution *y* = *y*(*x*, *γ*) exists, we write

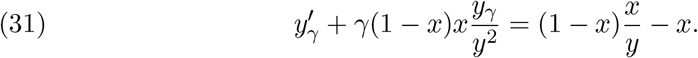

We see that the integrating factor

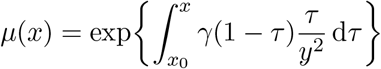

satisfies *μ*(*x*) ≥ 1 and is increasing. We get, after integrating,

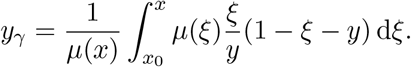

Since *μ*(*x*) ≥ 1 and the integrand is non-negative, we have *y_γ_* ≥ 0.

We now go one step further to investigate the sign of *y_γγ_*. Differentiating (31) with respect to *γ* we get

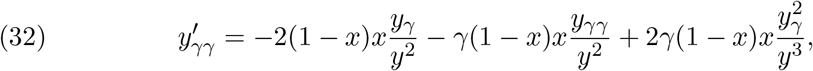

which is equivalent to

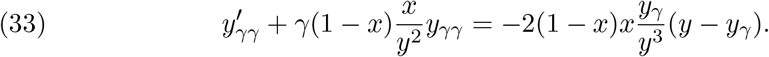

Multiplying by the same integrating factor *μ*(*x*), and integrating we see that *y_γγ_* is non-positive if *y* − *y_γ_* ≥ 0.

We see directly that *locally* we have *y_γγ_* ≤ 0, since *y_γ_*(*x*_0_) = 0 (the starting point does not depend on *γ*).

In forward time (when *y* = *y*(*x*) is further away from the initial value *y*_0_ = *y*(*x*_0_)) it is also true under some circumstances, *i.e.,* conditions on *x* and *y*, and possibly it is simply true for all 0 ≤ *x,y* ≤ 1. We will not go through all possible such conditions on *y* and *x* here, but as an example, suppose that *y* ≥ *x*. Then, since *μ*(*x*) is increasing, by the Mean Value Theorem for integrals, for some 0 ≤ *η* ≤ *x*,

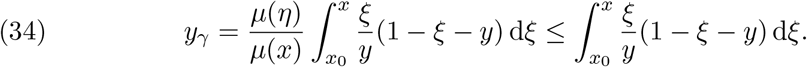

Using that *ξ* ≤ *y* we get

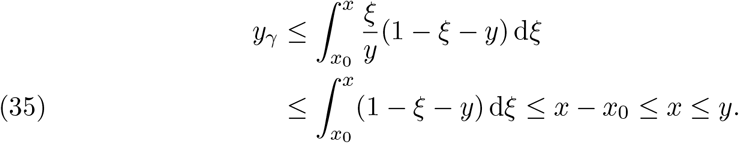

So as long as *y* ≥ *x* the statement holds that *y_γγ_* ≤ 0 (with strict inequality if *x,y* ≠ 0,1). If we require that *y* ≥ min(*x*, 1 − *x*), which is a slightly stronger condition, then, using that *y* ≥ *x* on [0,1/2] and that *y* ≥ 1 − *x* on [1/2,1],

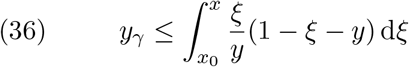

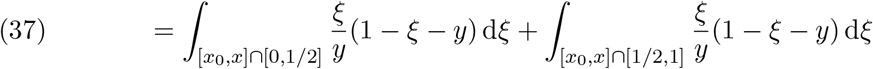

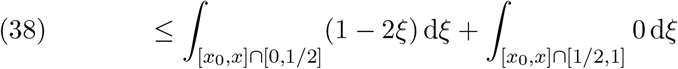

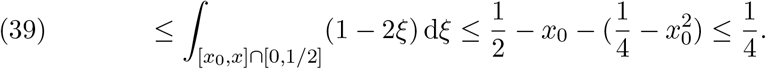

Combining this with the estimate just before (see (35)), this means that *y_γ_* ≤ *y* in the almost triangle shaped domain 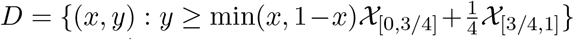, where *χ_I_* is the characteristic function on *I* (*i.e.*, the function that takes the value 1 on *I* and 0 elsewhere). Although it is very likely that these estimates can be improved, they show that the phenomenon we proved in the metacommunity model also holds in this simple compartment model, at least for a large proportion of the states in the phase space. Finally, we recall that the phenomenon holds locally around every initial point. This does not imply a global statement, since far away from these initial values, *x* = *R* and *y* = *S_R_* may take different routes for different parameters *a* and *β*.

## Supplement B: Statistical analysis of resistance data

### 1. Introduction

Here we analyse data on antibiotic resistance, antibiotic use, and consumption of alcohol-based hand rub in long-term care facilities (LTCF) given in the report “Point prevalence survey of healthcare-associated infections and antimicrobial use in European long-term care facilities. April-May 2013” from the European Centre for Disease Prevention and Control (ECDC), henceforth “the LTCF report” (25). We also use data on nation-wide resistance from the ECDC surveillance report ‘‘Antimicrobial resistance surveillance in Europe 2014” (26). The analyses were performed in R version 3.5.0 (21).

### 2. Description of the data

The dataset analysed includes for LTCF in each country the median consumption of alcohol-based hand rub in litres per 1000 resident days, the percentage of residents that were being treated with antibiotics at the time point when the study was performed, the percentage of treated residents receiving penicillins, the percentage receiving other *β*-lactams, the proportion of *Enterobacteriaceae* isolates from the facilities that were resistant to third generation cephalosporins, and the number of *Enterobacteriaceae* isolates that were tested for resistance. It also includes for each country the proportion of *E. coli* isolates that were resistant to third generation cephalosporins in nation-wide surveillance (not only LTCF) in the same year. The hand rub data (”Handrub”) were extracted from the map in Figure 17 in the LTCF report, and were coded as the centre of the given interval. Data given as ≥ 8 were coded as 10. The percentage of residents on antibiotic treatment was taken as the medians from Table 17 in the LTCF report, and the percentages of treated residents on different antibiotics (penicillins and other *β*-lactams, respectively) were taken from Table 19 in the LTCF report. The data on *Enterobacteriaceae* resistant to third generation cephalosporins were extracted by measurement in Figure 36 in the LTCF report, using Adobe Photoshop, and proportions were calculated for the isolates for which the resistance status was known. These proportions were then multiplied with the number of isolates tested to yield the counts of resistant and sensitive isolates. The proportion of *E. coli* resistant to third generation cephalosporins in nation-wide surveillance pertain to 2013, and were taken from Table 3.4 in the ECDC report ‘‘Antimicrobial resistance surveillance in Europe 2014”. Only countries that provided data for all variables were included in the analysis.

### 3. Statistical analyses

The log-odds of resistance to third generation cephalosporins in *Enterobacteriaceae* was modelled using logistic regression on the counts of resistant and sensitive isolates for each country. Two models were evaluated.

#### Model 1

The first model is that this (log-odds) of resistance, let us call it *L*, is

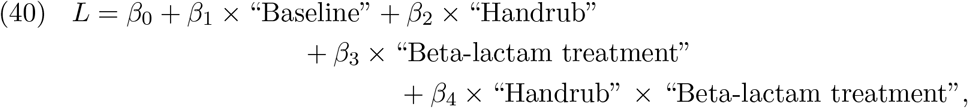

where “Baseline” is the proportion of *E. coli* resistant to third generation cephalosporins in each respective country as a whole, as described above, and “Beta-lactam treatment” is the prevalence of *β*-lactam (penicillins + other *β*-lactams) treatment in the LTCF in each country. The inclusion of the baseline means that the model estimates the effect of the use of hand rub and *β*-lactams in LCTFs as being proportional to the resistance in the corresponding country. Were we to estimate the difference in log-odds, we would need to put the *β*_1_ exactly equal to 1 in front of the resistance in each country as this would yield a difference between LTCFs and the country as a whole. This model posits that

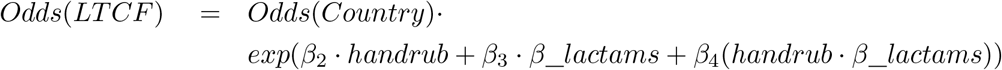

This is entirely feasible using an offset in the statistical software R. However, there is no strong prior reason to restrain *β*_1_ to be exactly equal to 1. Thus, we use the following model:

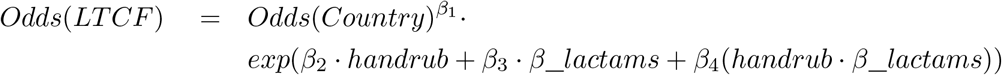

and retain this model if it gives reasonable output. The deviance table for this model is:

| | Df | Deviance | Resid-Df | Resid-Dev | Pr(> $\chi$) |
| --- | --- | --- | --- | --- | --- |
| NULL |  |  | 11 | 35.930 |  |
| Country | 1 | 15.9072 | 10 | 20.023 | $6.652 \cdot 10^{-5}$ *** |
| Handrub | 1 | 4.0251 | 9 | 15.997 | 0.044827 * |
| Beta-lactam-treatment | 1 | 3.1580 | 8 | 12.839 | 0.07557 |
| Handrub ×<br>Beta-lactam-treatment | 1 | 7.8964 | 7 | 4.943 | 0.004953 ** |

And the table of estimates is:

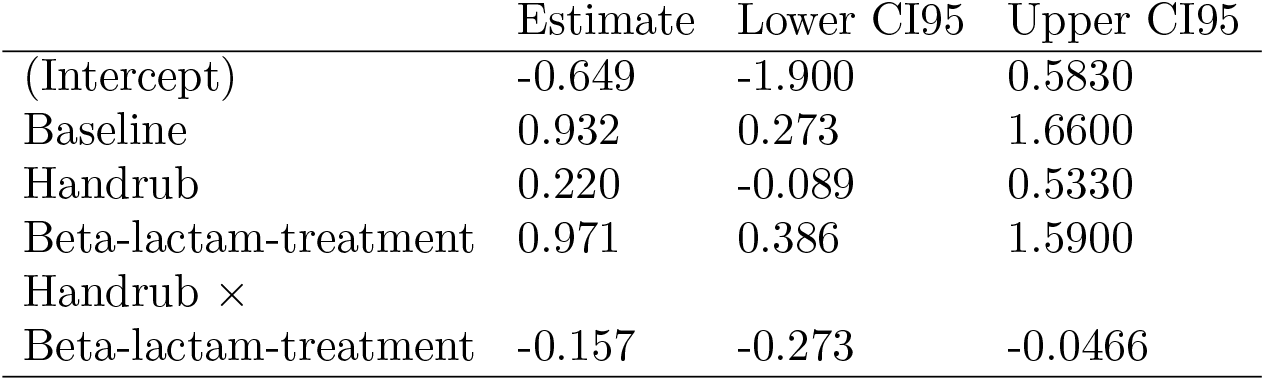

There is thus a significant negative interaction between “Handrub” and “Betalactam treatment”. The *p*-value for the interaction term is well below 0.05, and its CI95 does not include 0. This model has 5 degrees of freedom and an AIC of 43.15532.

#### Model 2

Since it is biologically unclear whether all *β*-lactams or only non-penicillin *β*-lactams should be included, we also use a model with only non-penicillin *β*-lactams. This model is (with *L* as above):

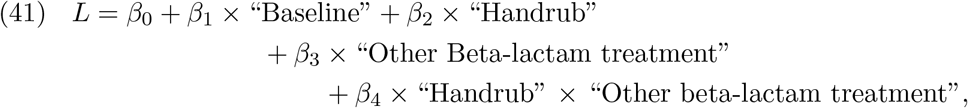

where “Other beta-lactam treatment” includes only non-penicillin *β*-lactams, and the other variables are defined as above. This model has 5 degrees of freedom and an AIC of 49.56753.

#### Model choice

We retain model 1, as it has the lower AIC.

#### Comments

The dataset has few observations, so if we were to consider confounding factors, a non-significant effect would not be evidence of lack of influence. Since the data are from an observational study, it is impossible to state that the model is correct. However, we believe that it contains the minimal set of explanatory variables.

## References

[1] B. G. Bell, F. Schellevis, E. Stobberingh, H. Goossens, and M. Pringle, “A systematic review and meta-analysis of the effects of antibiotic consumption on antibiotic resistance,” BMC Infectious Diseases, vol. 14, p. 13, Jan 2014.

[2] C. Costelloe, C. Metcalfe, A. Lovering, D. Mant, and A. D. Hay, “Effect of antibiotic prescribing in primary care on antimicrobial resistance in individual patients: systematic review and meta-analysis,” BMJ, vol. 340, 2010.

[3] L. M. King, K. E. Fleming-Dutra, and L. A. Hicks, “Advances in optimizing the prescription of antibiotics in outpatient settings,” BMJ, vol. 363, 2018.

[4] M. Baym, L. K. Stone, and R. Kishony, “Multidrug evolutionary strategies to reverse antibiotic resistance,” Science, vol. 351, no. 6268, 2016.

[5] D. McAdams, K. Wollein Waldetoft, C. Tedijanto, M. Lipsitch, and S. P. Brown, “Resistance diagnostics as a public health tool to combat antibiotic resistance: A model-based evaluation,” bioRxiv, 2018.

[6] C. Tedijanto, S. W. Olesen, Y. H. Grad, and M. Lipsitch, “Estimating the proportion of bystander selection for antibiotic resistance among potentially pathogenic bacterial flora,” Proceedings of the National Academy of Sciences, vol. 115, no. 51, pp. E11988–E11995, 2018.

[7] D. Pittet and J. M. Boyce, “Hand hygiene and patient care: pursuing the Sem-melweis legacy,” The Lancet Infectious Diseases, vol. 1, pp. 9–20, 2001. Preview Issue.

[8] WHO, SAVE LIVES: Clean Your Hands 5 May 2017: Fight antibiotic resistance-its in your hands. 2017.

[9] J. O’Neill, “Infection prevention, control and surveillance: limiting the development and spread of drug resistance,” The Review on Antimicrobial Resistance, March 2016.

[10] M. A. Leibold and J. M. Chase, Metacommunity Ecology (MPB-59). Princeton University Press, 2018.

[11] J. R. Mihaljevic, “Linking metacommunity theory and symbiont evolutionary ecology,” Trends in Ecology & Evolution, vol. 27, no. 6, pp. 323–329, 2012.

[12] F. Blanquart, “Evolutionary epidemiology models to predict the dynamics of antibiotic resistance,” Evolutionary Applications (doi:10.1111/eva.12753).

[13] M. Vellend, The theory of ecological communities (MPB-57), vol. 75. Princeton University Press, 2016.

[14] T. Day, S. Huijben, and A. F. Read, “Is selection relevant in the evolutionary emergence of drug resistance?,” Trends in Microbiology, vol. 23, no. 3, pp. 126–133, 2015.

[15] E. Hansen, R. J. Woods, and A. F. Read, “How to use a chemotherapeutic agent when resistance to it threatens the patient,” PLOS Biology, vol. 15, pp. 1–21, 02 2017.

[16] A. R. Wargo, S. Huijben, J. C. de Roode, J. Shepherd, and A. F. Read, “Competitive release and facilitation of drug-resistant parasites after therapeutic chemotherapy in a rodent malaria model,” Proceedings of the National Academy of Sciences, vol. 104, no. 50, pp. 19914–19919, 2007.

[17] T. Day and A. F. Read, “Does high-dose antimicrobial chemotherapy prevent the evolution of resistance?,” PLOS Computational Biology, vol. 12, pp. 1–20, 01 2016.

[18] S. Huijben, A. S. Bell, D. G. Sim, D. Tomasello, N. Mideo, T. Day, and A. F. Read, “Aggressive chemotherapy and the selection of drug resistant pathogens,” PLOS Pathogens, vol. 9, pp. 1–9, 09 2013.

[19] S. P. Hubbell, “A unified theory of biogeography and relative species abundance and its application to tropical rain forests and coral reefs,” Coral Reefs, vol. 16, pp. S9–S21, Jun 1997.

[20] S. P. Hubbell, The unified neutral theory of biodiversity and biogeography (MPB-32). Princeton University Press, 2001.

[21] R. Core Team, “R: A language and environment for statistical computing,” R Foundation for Statistical Computing, Vienna, Austria. URL https://www.R-project.org/., 2018.

[22] N. Bruinsma, J. Hutchinson, A. van den Bogaard, H. Giamarellou, J. Degener, and E. Stobberingh, “Influence of population density on antibiotic resistance,” Journal of Antimicrobial Chemotherapy, vol. 51, pp. 385–390, 2 2003.

[23] WHO, WHO Guidelines on Hand Hygiene in Health Care: First Global Patient Safety Challenge: Clean Care is Safer Care. 2009.

[24] V. Erasmus, T. J. Daha, H. Brug, J. H. Richardus, M. D. Behrendt, M. C. Vos, and E. F. van Beeck, “Systematic review of studies on compliance with hand hygiene guidelines in hospital care,” Infection Control and Hospital Epidemiology, vol. 31, no. 3, p. 283–294, 2010.

[25] European Centre for Disease Prevention and Control (ECDC), Point prevalence survey of healthcare-associated infections and antimicrobial use in European long-term care facilities. April-May 2013. 2014.

[26] European Centre for Disease Prevention and Control (ECDC), Antimicrobial resistance surveillance in Europe 2014. 2015.

